# Predictive Modeling of Gene Expression and Localization of DNA Binding Site Using Deep Convolutional Neural Networks

**DOI:** 10.1101/2024.12.17.629042

**Authors:** Arman Karshenas, Tom Röschinger, Hernan G. Garcia

**Affiliations:** Biophysics Graduate Group, University of California at Berkeley, Berkeley, CA, USA; Division of Biology and Biological Engineering, California Institute of Technology, Pasadena, CA, USA; Department of Molecular and Cell Biology, University of California Berkeley, Berkeley, CA, USA; Department of Physics, University of California, Berkeley, CA, USA; Institute for Quantitative Biosciences-QB3, University of California, Berkeley, CA, USA; Chan Zuckerberg Biohub – San Francisco, San Francisco, CA, USA

## Abstract

Despite the sequencing revolution, large swaths of the genomes sequenced to date lack any information about the arrangement of transcription factor binding sites on regulatory DNA. Massively Parallel Reporter Assays (MPRAs) have the potential to dramatically accelerate our genomic annotations by making it possible to measure the gene expression levels driven by thousands of mutational variants of a regulatory region. However, the interpretation of such data often assumes that each base pair in a regulatory sequence contributes independently to gene expression. To enable the analysis of this data in a manner that accounts for possible correlations between distant bases along a regulatory sequence, we developed the Deep learning Adaptable Regulatory Sequence Identifier (DARSI). This convolutional neural network leverages MPRA data to predict gene expression levels directly from raw regulatory DNA sequences. By harnessing this predictive capacity, DARSI systematically identifies transcription factor binding sites within regulatory regions at single-base pair resolution. To validate its predictions, we benchmarked DARSI against curated databases, confirming its accuracy in predicting transcription factor binding sites. Additionally, DARSI predicted novel unmapped binding sites, paving the way for future experimental efforts to confirm the existence of these binding sites and to identify the transcription factors that target those sites. Thus, by automating and improving the annotation of regulatory regions, DARSI generates experimentally actionable predictions that can feed iterations of the theory-experiment cycle aimed at reaching a predictive understanding of transcriptional control.

## Introduction

A central challenge in biology is to accurately predict gene regulatory programs and their functions from knowledge of genome sequences (***Pennacchio et al., 2013***; ***Phillips et al., 2019***; ***Bintu et al., 2005***). These programs are governed, in large part, by DNA regulatory regions containing binding sites for transcription factors. These proteins interact with the transcriptional machinery to modulate gene expression by enhancing or repressing transcription.

Achieving such predictive understanding of transcriptional regulation requires addressing two key challenges: (i) identifying and characterizing transcription factor binding sites within regulatory regions and (ii) integrating this knowledge into theoretical models capable of quantitatively predicting how the number, placement and affinity of these binding sites dictate gene expression (***Stormo, 2000***; ***Phillips et al., 2019***; ***Bintu et al., 2005***). Thus, the foundational step toward predicting the regulatory outcomes encoded by DNA regulatory regions involves determining the location and identity of transcription factor binding sites.

Despite the key need to map transcription factor binding sites in regulatory regions, our ability to accurately identify these sites is still lacking (***Minchin and Busby, 2009***; ***Santos-Zavaleta et al., 2018***). For instance, in the bacterium *Escherichia coli*, one of the most thoroughly studied model organisms, binding sites regulating only about 33% of genes have been mapped to date (***Tierrafría et al., 2022***; ***Ireland et al., 2020***; ***Santos-Zavaleta et al., 2018***). While some genes may not be transcriptionally regulated and thus lack transcription factor binding sites, this figure more likely reflects the limited number of detailed case studies conducted so far. The challenge is even more pronounced in multicellular organisms, such as the fruit fly *Drosophila melanogaster*, where regulatory networks are considerably more intricate and less well characterized (***Keränen et al., 2022***).

Classic approaches for finding and validating binding sites within regulatory regions are typically manual and, therefore, low-throughput. Specifically, these approaches rely on the creation of reporter constructs where suspected binding sites are mutagenized. By correlating DNA sequence with the resulting reporter expression level, transcription factor binding sites can be validated. As a result of the low-throughput nature of this pipeline, the binding sites controlling only a handful of genes in model organisms have been mapped in detail (e.g., ***Müller-Hill (1996***); ***Schleif (2003***); ***Ptashne (2004***); ***Weickert and Adhya (1993***); ***Levine (2010***)).

Massively Parallel Reporter Assays (MPRAs) have recently emerged as a powerful tool for mapping regulatory sequences (***Patwardhan et al., 2009***; ***Kinney et al., 2010***; ***Patwardhan et al., 2012***; ***Melnikov et al., 2012***; ***Kwasnieski et al., 2012***; ***Kreimer et al., 2022***; ***Zheng and VanDusen, 2023***; ***Ireland et al., 2020***; ***Belliveau et al., 2018***). These assays involve synthesizing a large library (>1,000s) of mutagenized variants of a regulatory region and incorporating them into plasmids (Fig. 1A,B). The plasmid library is then transfected into cells, where, after cell lysis, gene expression levels for each variant are measured in high-throughput using sequencing (Fig. 1C,D).

**Figure 1.**
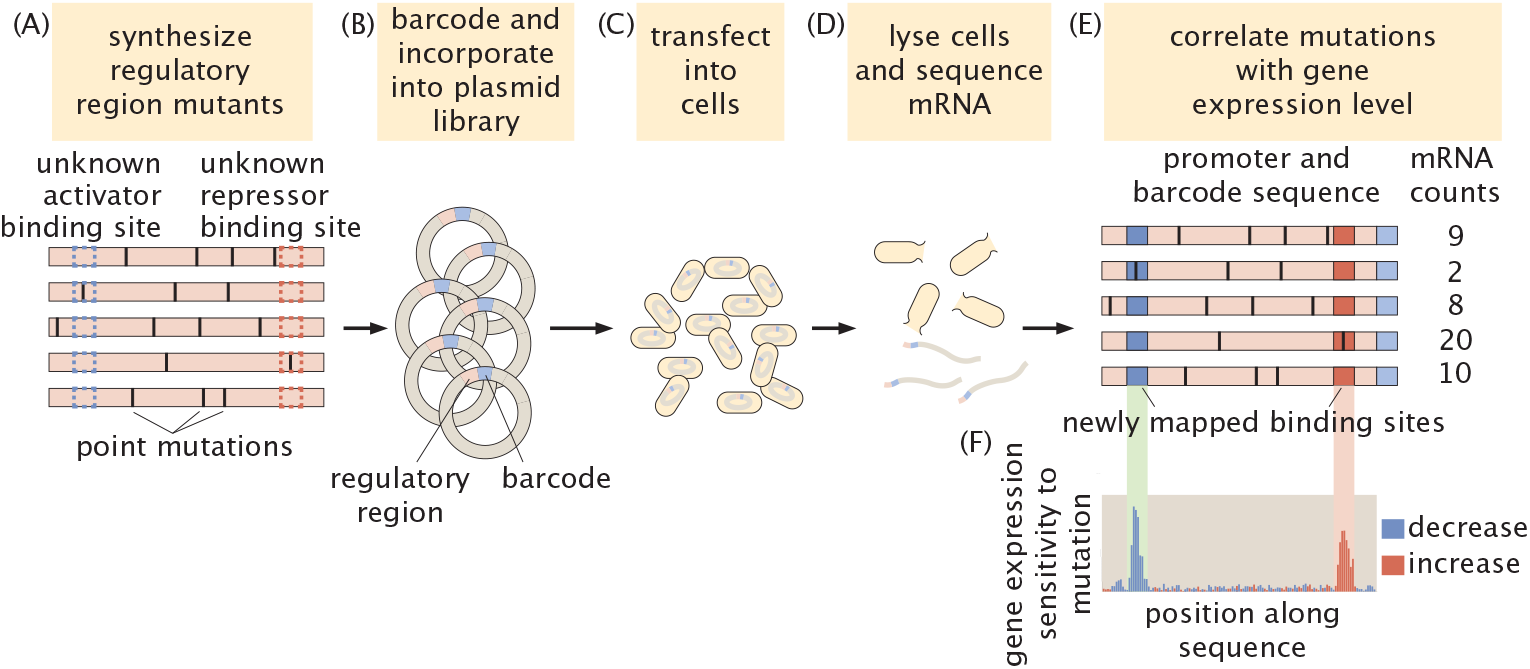
Schematic example of a massively parallel reporter assay to dissect regulatory regions in *E. coli*. **(A)** A library of mutated versions of a previously uncharted regulatory sequence is synthesized. **(B)** Each sequence is barcoded and incorporated into constructs that drive the expression of a reporter gene, forming a plasmid library. **(C)** The plasmid library is transformed into cultured cells such as *E. coli*. **(D)** After cell lysis, the reporter mRNA is extracted and quantified by sequencing. **(E)** An illustrative example showing the correlation between the regulatory sequence variants and their corresponding gene expression levels. **(F)** These data make it possible to capture the shift in gene expression upon mutagenesis of each base pair along the sequence, leading to the identification of activator and repressor binding sites.

By linking the sequences of these mutated regulatory regions to their corresponding gene expression levels (Fig.1E), MPRAs allow for the identification of positions within the sequence that influence gene expression when mutated. As illustrated in Figure 1F, this approach makes it possible to pinpoint transcription factor binding sites in uncharacterized regulatory regions: mutations in activator binding sites typically decrease gene expression, whereas mutations in repressor binding sites tend to increase expression (***Ireland et al., 2020***; ***Belliveau et al., 2018***).

While MPRAs have significantly advanced the study of regulatory sequences (***Kreimer et al., 2022***; ***Zheng and VanDusen, 2023***), key challenges remain in systematically analyzing the resulting datasets to reveal transcription factor binding sites. For example, an important potential limitation lies in the reliance of these analyses on metrics such as gene expression sensitivity to mutation (Fig. 1F) or mutual information between gene expression and base pair identity (***Kinney et al., 2010***; ***Ireland et al., 2020***). These measures often assume that base pairs contribute independently to gene expression: because these metrics evaluate the impact of mutations at specific positions by effectively averaging their effects across all other positions in the sequence, they potentially ignore nucleotide interactions within the regulatory sequence.

In this study, we address the challenges of finding binding sites and developing predictive models in a manner that can account for potential spatial correlations along DNA sequences by introducing a novel computational framework, the Deep Learning Adaptive Regulatory Sequence Identifier (DARSI). DARSI capitalizes on recent advancements in convolutional neural networks and deep learning (***Alzubaidi et al., 2021***; ***LeCun et al., 2015***; ***Park and Kellis, 2015***; ***de Almeida et al., 2022***; ***Avsec et al., 2021a***; ***Kelley et al., 2018***; ***Zrimec et al., 2021***) to capture nucleotide interactions distributed throughout the sequences assayed by MPRAs. DARSI makes it possible to predict gene expression levels—albeit these levels are discretized—from raw regulatory sequences without relying on prior knowledge of the underlying regulatory architecture.

The *predictive power* enabled by DARSI, although far from the *predictive understanding* we ultimately seek through physical models (***Barnes et al., 2019***; ***Kinney et al., 2010***; ***Belliveau et al., 2018***; ***Tareen et al., 2022***; ***Pan et al., 2024***; ***Lagator et al., 2022***), makes it possible to obtain detailed insights into the number and spatial arrangement of transcription factor binding sites within regulatory sequences. Hypothesized binding sites are identified through the integration of saliency mapping techniques—akin to an *in silico* mutagenesis experiment—which allow us to interpret the impact of specific nucleotide sequence changes on gene expression outcomes.

We applied DARSI to MPRA data from 95 operons in *E. coli* published by ***Ireland et al. (2020***). First, we demonstrate that the trained networks achieve an average accuracy of ∼80% in predicting the expression levels of the reporter gene directly from the raw sequences. Building on this predictive power, we show that the networks can be leveraged to identify transcription factor binding sites. Specifically, DARSI identified over 170 binding sites, including more than 88% of the previously mapped sites (***Tierrafría et al., 2022***), and uncovered 73 new hypothesized binding sites across these operons. Thus, we demonstrate that the convolutional neural network architecture within DARSI can be used to augment analyses of gene expression MRPA data to both achieve predictive power and identify binding sites that can guide further experiments.

## Results

### DARSI: A Convolutional Neural Network for Gene Expression Prediction from MPRA Data

To reach predictive power over regulatory regions and capture correlations between nucleotides from MPRA data, we developed a convolutional neural network. The convolutional filters within this network are capable of modeling interactions between distant base pairs (***Alipanahi et al., 2015***), potentially making it possible to identify regulatory features that span different regions of the DNA sequence. The network takes as input sequence variants of a given operon and corresponding expression levels of the reporter gene. The architecture we converged on after optimization (see “DARSI Architecture and Training” section of the Supplementary Information) consists of 12 layers and is similar to previously established models in the field (***Avsec et al., 2021a***,b; ***Alipanahi et al., 2015***).

As a case study, we utilized data from a recent MPRA study conducted by ***Ireland et al. (2020***) in *E. coli*. This work dissected the regulatory information of 114 bacterial operons by randomly mutating a 160 bp region upstream of the transcription start site of each operon at a mutation rate of 10%. This process generated a dataset akin to that featured in the schematic shown in Figure 2A that correlates sequence and gene expression. To ensure sufficient coverage of mutations, we selected operons with at least 1,000 sequence variants. This number guaranteed that, for every base pair along the sequence, our dataset contained at least 100 sequence variants in which that base pair is mutated. We further cross-referenced all sequences with the annotated *E. coli* genome available on *EcoCyc* (***Moore et al., 2024***) to verify that the sequences encompassed regions upstream of the genes. The lower bound used for number of variants and the cross-validation of the data with annotated databases reduced the dataset to 95 mutagenized operons, each originating from *E. coli* colonies cultivated in LB medium. Across the 95 operons, the mean number of unique sequences per operon is 2083 ±960, with 847 ±193 unique barcodes per operon. This results in an overall mean of 8313 ± 3228 sequence variants across all operons. The sequence data for each operon served as input to the network, while discretized normalized mRNA counts (described below) were used as the output. A separate convolutional neural network was trained for each operon, resulting in a total of 95 independently trained networks.

**Figure 2.**
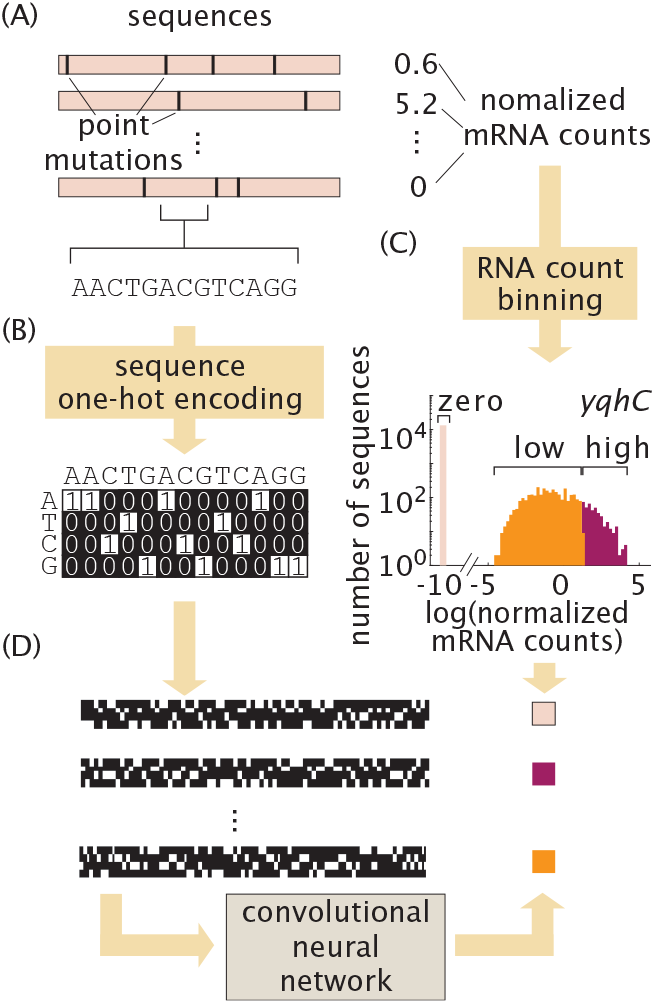
The DARSI pipeline. **(A)** MPRA dataset that makes it possible to correlate regulatory sequence with gene expression. **(B)** One-hot encoding scheme used to convert each DNA sequence into a binary image. **(C)** Distribution of the log(normalized mRNA count) together with the bin of gene expression assigned to each value for the illustrative case of the *yqhC* operon. Note that the overlap observed between the low and high expression bins is an artifact of the histogram binning and does not reflect an actual overlap between the classes of expression. **(D)** The images and the corresponding gene expression bins constitute the inputs and outputs of our convolutional neural network, respectively.

Convolutional neural networks are designed to take images or matrices as inputs. Thus, to prepare the DNA sequence data for use as input to our networks, we transformed the sequences into a two-dimensional matrix representation. Specifically, each 160 bp regulatory sequence from the MPRA dataset was encoded as a 4×160 binary image using a so-called one-hot encoding scheme, as illustrated in Figure 2B and detailed in the “One-Hot Encoding” section of the Materials and Methods. Consequently, the data for each operon is represented as a stack of 4 × 160 images, with each image corresponding to a specific sequence variant for that operon.

As output from the network, we separated gene expression levels into discrete bins. We then used the networks to predict which gene expression bin regulatory sequences correspond to. Before discretization, we first normalized mRNA counts by dividing the number of sequenced mRNAs by the copy number of each regulatory sequence reported by DNA sequencing of the library (Fig. 2A). The objective of this normalization is to account for the fact that different regulatory sequences will be present at different copy numbers in the library.

To train a convolutional neural network to classify sequences based on their corresponding gene expression levels, we categorized the normalized expression counts into discrete bins, referred to as “classes”. Mutations within the sequences can lead to various outcomes, such as a complete absence of detectable gene expression, a measurable reduction, or an increase in gene expression compared to typical levels observed across the sequence variants for each operon. To capture this variation in expression level, we examined the distribution of the logarithm of the normalized expression counts. Based on this distribution, we defined three distinct expression classes: (1) sequences resulting in no detectable gene expression (zero expression bin), (2) sequences yielding low but measurable levels of gene expression (low expression bin), and (3) sequences associated with high levels of gene expression (high expression bin). While the decision to use three bins was informed by the natural clustering of data in the logarithmic space, this choice is ultimately a simplification that, as we will show in the next sections, can still lead to predictive power and the ability to identify transcription factor binding sites.

For each operon we determined the thresholds of log(normalized mRNA count) for each bin to partition the gene expression counts into the three classes. The zero gene expression bin corresponds to sequences that yielded no detectable mRNA. The threshold between the low and high gene expression bins were chosen so as to lead to statistically significant differences in mean gene expression levels between these two classes, as described in detail in the “RNA count labeling” section of the Materials and Methods. Figure 2C shows the distribution of log(normalized mRNA count) and the associated bins color-coded for the illustrative *yqhC* operon from the MPRA dataset by ***Ireland et al. (2020***). This operon is used throughout the text to illustrate our pipeline and its results, as it represents the average performance of the pipeline. Similar plots to Figure 2C, showing the distribution of expression counts for the rest of the operons in the dataset can be accessed through our GitHub repository.

The number of observations in each expression bin vary significantly. Indeed, as shown in Figure 2C, the bin corresponding to zero gene expression was typically overrepresented with respect to the low and high gene expression bins. To account for this over-representation, the zero gene expression bin was under sampled when training the networks, while the low and high bins were over sampled to create an evenly split processed dataset (***Bowyer et al., 2011***; ***He and Garcia, 2009***).

Training for each network utilized 70% of the processed data, following adjustments for data imbalance. Training was conducted in *MATLAB* using standard optimization toolboxes, with parameters optimized via stochastic gradient descent (***MathWorks, 2022***; ***Bottou, 1998***; ***Sra et al., 2011***). An additional 15% of the data (3,000–5,000 variants across the 95 operons) was reserved for validation during training, serving to optimize network architecture as discussed below. The remaining 15% of the data was allocated for final evaluation of the predictive power of each network.

Before engaging in the training of all 95 networks, we optimized the overall network architecture for accuracy in predicting gene expression in our dataset. While adding more convolutional layers should allow the network to extract longer-range interactions between nucleotides along the sequence, increasing the depth of the network leads to a substantial rise in the number of trainable parameters, potentially resulting in overfitting (***Alzubaidi et al., 2021***). As a result, we systematically and iteratively modulated the network architecture to assess its impact on prediction accuracy.

To optimize the network architecture, we focused on data from the 10 operons with the largest number of sequence variants. As expected, our optimization revealed that increasing model complexity (e.g., by adding layers and channels) generally improves training accuracy but can lead to overfitting, where the model performs poorly when validated using unseen data (Fig. S1). We converged onto an optimal architecture, detailed in Table S2, that strikes a balance between model complexity and performance. This chosen architecture is consistent with similar networks implemented in prior studies (***Avsec et al., 2021a***,b; ***de Almeida et al., 2022***). Using this optimized architecture, we independently trained 95 convolutional neural networks, one for each operon in our dataset. Further details on the DARSI architecture, its optimization, and training specifications can be found in the “DARSI Architecture and Training” section of the Supplementary Information.

### DARSI Can Predict Gene Expression from Raw Sequence

As outlined above, each trained network, corresponding to an individual operon in the dataset, was evaluated using the reserved 15% of the processed data designated as the test set. For each operon, raw sequences from the test partition were input into the trained network, which then predicted the corresponding output gene expression bin. The predicted bins were compared against the experimentally measured gene expression values to calculate an accuracy score for each network. As illustrated in Figure 3A, the networks achieved an average predictive accuracy of 79.8% across all 95 operons.

**Figure 3.**
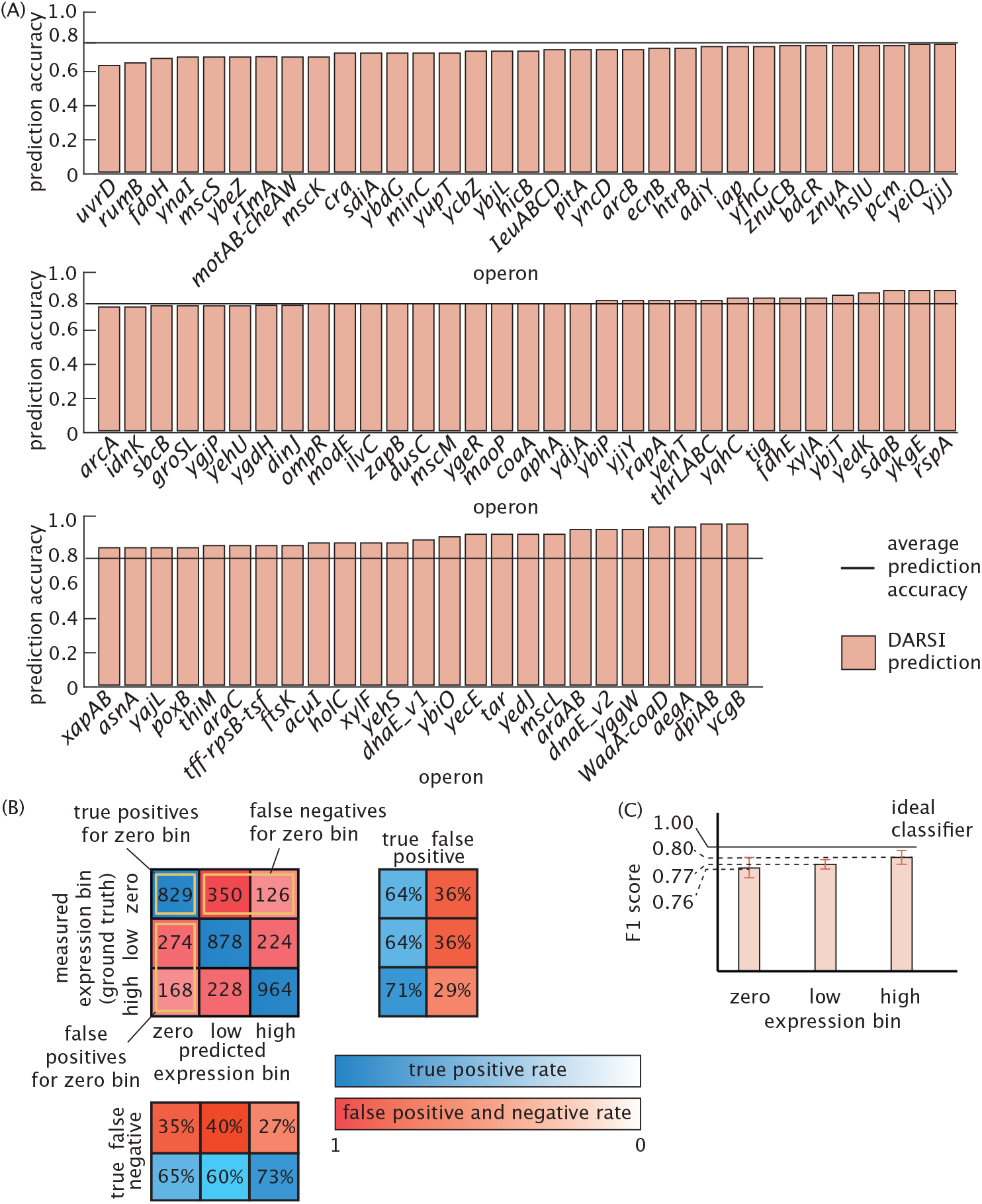
Predictive Performance of DARSI Models on Bacterial MPRA Data. **(A)** The accuracy of the DARSI model in predicting gene expression levels from previously unseen DNA sequences for each *E. coli* operon in the dataset. The average prediction accuracy of 79.8% is shown as a horizontal line. **(B)** Confusion matrix illustrates the comparison between measured and predicted expression levels for the *yqhC* operon in *E. coli*, highlighting model performance. Each entry in the matrix represents the number of sequences classified as true positives, true negatives, false positives and false negatives for each gene expression bin. The row projections of the confusion matrix in blue and red are true positive and false positive rates, respectively, while the column projections in blue and red are the true negative and false negative rates, respectively. **(C)** Average F1 score values for the three expression bins are shown with the ideal classifier represented by the solid horizontal line. The values of the F1 score are close to 1, corresponding to an ideal classifier.

To more rigorously evaluate the effectiveness of our DARSI model in predicting gene expression from raw sequence input, we generated confusion matrices. In these matrices, each column represents predicted expression bin (i.e., zero expression, low expression or high expression), while each row indicates the actual bin to which the sequences belong as reported by measurements. Each entry within the matrix indicates the number of sequences belonging to each combination of predicted and measured gene expression bins. Consequently, these matrices provide a summary of false positives, false negatives, true positives, and true negatives for each of the three discrete expression bins.

The confusion matrix for the *yqhC* operon is displayed in Figure 3B. This matrix indicates that the trained model classifies the majority of unseen data for each operon with *high specificity* (low false positive rate) and *high sensitivity* (low false negative rate), as evidenced by the diagonal dominance and the row and column projections shown in Figure 3B. To access a full list of confusion matrices for all the models trained, the reader is referred to the GitHub repository.

To evaluate the the overall performance of DARSI across all 95 operons, we computed the average F1 score for each expression bin. The F1 score is a metric that assesses both specificity and sensitivity of classifiers, and is commonly employed to gauge classifier performance (***Sokolova and Lapalme, 2009***; ***Powers, 2020***; ***Alzubaidi et al., 2021***; ***Aloysius and Geetha, 2017***). The F1 score for a given bin of expression is defined as

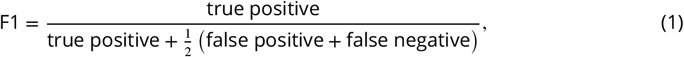

where, for example, “true positive” indicates the number of true positives resulting from our model for a specific bin. According to this definition, an ideal classifier with 100% true positive and 0% false positive and false negative rates will have an F1 score of one. True positives, false negatives, and false positives have been highlighted for the zero gene expression bin of the representative *yqhC* operon in Figure 3B, leading to an average F1 score of 0.64 across the three bins for this operon.

By averaging the F1 score of all DARSI networks, we can compare the average network performance to that of an ideal classifier. Figure 3C presents the F1 score values for the zero expression bin (0.76 ± 0.10), low expression bin (0.77 ± 0.09) and high expression bin (0.80 ± 0.09) averaged over all 95 trained convolutional neural networks, where the error bars indicate the standard deviation. The F1 scores for all three expression bins exceed the threshold of 0.7 that is commonplace in most fields (***Lipton et al., 2014***; ***Hicks et al., 2022***), indicating that the model effectively distinguishes sequences within these bins with high specificity and sensitivity. Thus, we deemed the gene expression predictions made by the trained DARSI models to be reliable for them to be leveraged in our exploration of regulatory architectures in *E. coli*.

### Uncovering Binding Sites Using Saliency Maps

We next leveraged the predictive power of our networks to identify transcription factor binding sites within unmapped regulatory regions. To achieve this task using MPRA datasets, existing methods typically analyze the mutual information or the shift in gene expression resulting from mutations at individual base pairs (Fig. 1F; ***Ireland et al. (2020***); ***Pan et al. (2024***); ***Belliveau et al. (2018***); ***Kheradpour et al. (2013***)). These analyses aim to isolate the impact of mutations at each nucleotide on the overall gene expression levels. To make this possible, the contributions of mutations across all other nucleotides within the sequence are averaged. Consequently, these approaches assume that the contributions of individual base pairs to gene expression are independent from one another. In contrast, the convolutional neural network architecture employed in this study makes it possible to account for spatial correlations between nucleotides throughout the sequence.

To examine the capacity of our networks to identify clusters of nucleotides corresponding to binding sites, we created so-called saliency maps. These saliency maps are better understood in the context of convolutional neural networks for image classification. For example, a neural network can be trained to classify images between those featuring a dog and those not featuring a dog (***Russakovsky et al., 2015***; ***Selvaraju et al., 2017***; ***Vinogradova et al., 2020***). A saliency map is a heat map that reports on how important each pixel within an image was in making the decision of how to classify that image. In the specific case of the classification of images featuring a dog, we would expect pixels that fall within the dog to carry more information than those pixels that are in the background of the image.

Similarly, because regulatory sequences (Fig. 4A) are represented as images using the one-hot encoding approach (Fig. 4B), the saliency map of each sequence describes how important each pixel within the image—a binary pattern unique to each sequence—is for determining the gene expression bin corresponding to that sequence (Fig. 4C). As a result, these saliency maps can be loosely thought of as heatmaps reporting on the sensitivity of the predicted gene expression level to mutating each nucleotide at every position along the regulatory sequence. As described in detail in the “Saliency Maps” section of the Materials and Methods, the generation of these maps involves calculating the derivative of the network loss function—a measure of how well the network does at predicting output gene expression—with respect to each pixel of the images encoding for the DNA sequence in the binary input layer of the network.

**Figure 4.**
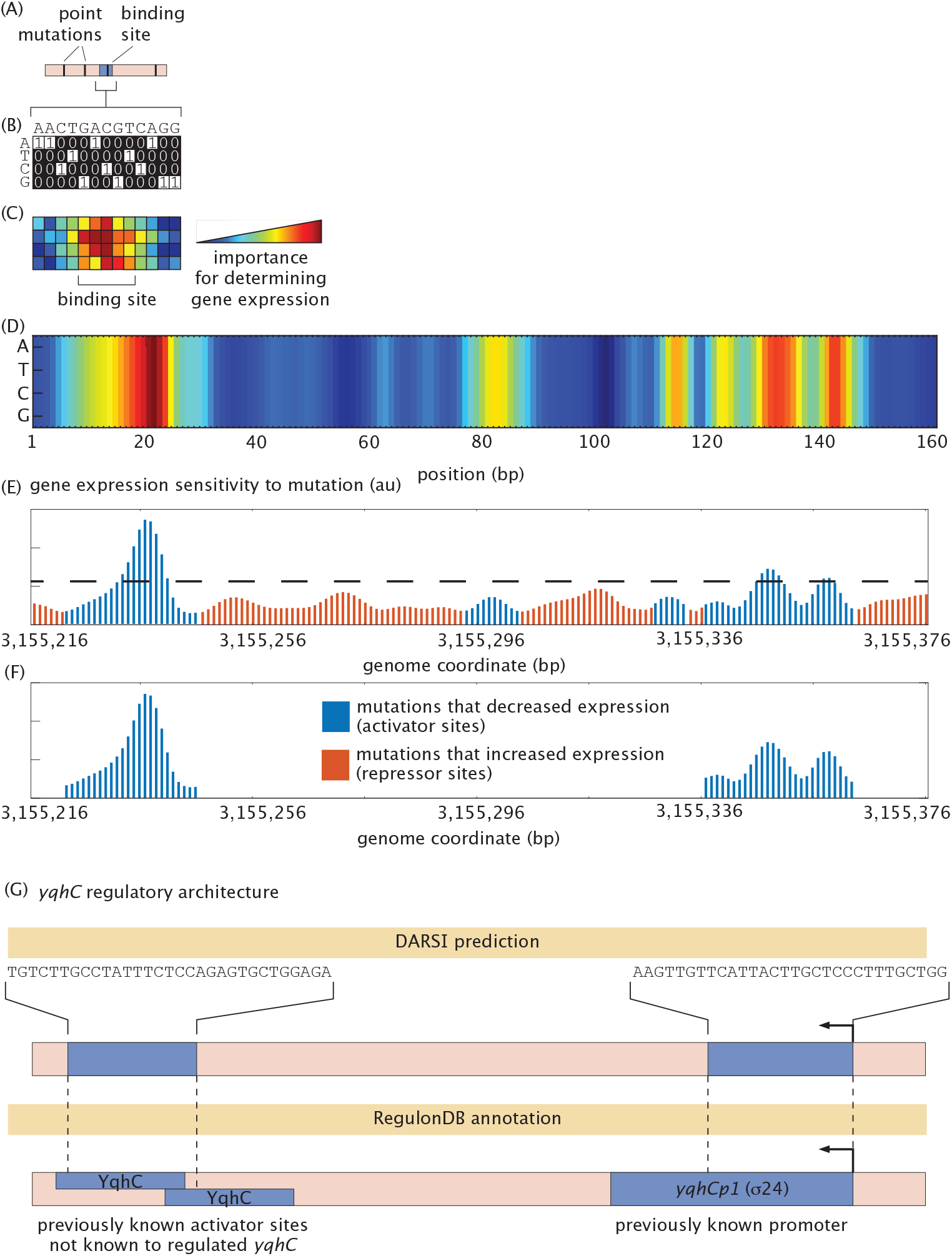
Saliency map generation and binding site identification. **(A)** Sequence variants from the test subset of a given operon, each containing random point mutations, are processed through the pre-processing pipeline to generate one-hot encodings, as illustrated in **(B). (C)** These one-hot encoded sequences are input into the trained DARSI model, where the gradient of the network loss is computed with respect to each pixel in the input image for each variant. These gradients measure the sensitivity of the network output to each nucleotide. Gradients are averaged across all variants in the test subset to generate saliency maps, which are represented as 4 × 160 heatmaps. These heatmaps indicate the pixels containing the most information used by the network to classify the gene expression bin of the input sequence. **(D)** An example saliency map for the illustrative *yqhC* operon highlights regions of high sensitivity. Notably, the saliency values at the same sequence position are relatively insensitive to base pair identity, suggesting that DARSI primarily relies on positional information rather than specific nucleotide identity to predict gene expression. **(E)** Maximum saliency values at each position are normalized and exponentiated to produce unitless plots of gene expression sensitivity to mutations, as shown for the *yqhC* operon. **(F)** These sensitivity plots are further refined by filtering peaks that exceed one standard deviation above the mean (dashed line in **(E)**), span at least 10 bp, and show contiguous effects as either activators or repressors. **(G)** The refined plots enable the identification of potential binding sites and operon regulatory architectures. For the *yqhC* operon, two previously annotated regions were identified: a promoter associated with the operon and an activator binding site, which, although annotated, had not been previously associated with the regulation of *yqhC*.

By applying this process to all sequence variants of a given operon in the test dataset, we generated an ensemble of saliency maps, one for each sequence variants. These maps are then averaged to produce a final saliency map for the operon. As shown in Figure 4D, the saliency map for the illustrative *yqhC* operon used throughout this study reveals several segments along the sequence that exhibit higher information content for predictions made by the trained DARSI model. Notably, minimal variation is observed along individual columns, suggesting that the network primarily considers the positional context of the base pair rather than its specific nucleotide identity when classifying expression levels. These clusters of highly sensitive positions form the initial hypotheses for the locations of binding sites within the sequence.

To interpret the information encoded within the saliency maps, the maximum saliency value among the four nucleotides at each position along the regulatory sequence—that is, along each column of the saliency map—is calculated. The result is a saliency vector that reports on the sensitivity of output gene expression to mutation along the regulatory sequence. Note that the absolute values of saliency maps generated by the network are not inherently interpretable; only relative changes in these values are meaningful. As a result, we normalize the saliency vector by subtracting its mean and dividing by its standard deviation. Subsequently, the normalized saliency vectors are exponentiated to represent likelihoods or probabilities (see the “Binding Site Identification” section in the Materials and Methods). These processed values are visualized as “gene expression sensitivity to mutation” plots. Figure 4E shows an example of this plot for the *yqhC* operon. Because we are after binding site-sized features within these plots, we smoothed the curve by averaging the data using a sliding window of size 5 bp as in previous studies (***Ireland et al., 2020***).

To identify transcription factor binding sites, we examined the smoothed values of the gene expression sensitivity to mutation shown in Figure 4E. Here, red bars correspond to positions along the sequence where mutations led to an increase in expression, suggesting potential repressor binding sites. Conversely, blue bars represent nucleotides where mutations resulted in decreased expression, indicating potential activator binding sites.

To predict binding sites along regulatory sequences, we identify clusters of base pairs with high sensitivity. Specifically, following the approach of ***Robison et al. (1998***), we detect positions where the maximum sensitivity exceeds one standard deviation above the mean sensitivity across the entire sequence (horizontal line in Fig. 4E). In accordance with the minimum length of DNA binding sites in *E. coli* reported by ***Stewart et al. (2012***), ***Rydenfelt et al. (2015***), and ***Ruths and Nakhleh (2013***), potential binding sites are defined as regions exceeding this threshold and spanning at least 10 base pairs. Figure 4F presents a filtered expression sensitivity-to-mutation plot, highlighting two prominent peaks (blue) corresponding to regulatory regions annotated in RegulonDB (***Tierrafría et al., 2022***). The first peak aligns with a promoter previously mapped to *yqhC*, while the second corresponds to activator binding sites that, although annotated in RegulonDB, had not been associated with the regulation of *yqhC*. This finding highlights DARSI’s ability to identify functional connections between regulatory elements and their target genes. Notably, these activator binding sites are not obvious when examining the mutual information (Fig. S4A), which constitutes the basis of previous approaches for identifying binding sites using MPRA data (***Ireland et al., 2020***). This particular example demonstrates DARSI’s capacity to reveal regulatory features overlooked by traditional methods. A detailed description of the filtering steps employed to generate these expression shift plots is provided in the “Binding Site Identification” section of the Materials and Methods. Further, a comparison of the sensitivity of all operons predicted by DARSI to the same analysis based on mutual information can be found in the GitHub repository.

Using this pipeline, we identified a total of 172 binding sites across all 95 operons, successfully capturing 88.4% of the previously documented sites in published and curated databases (***Tierrafría et al., 2022***). In addition to these annotated binding sites, DARSI predicted 73 hypothetical novel binding sites, spanning more than one-third of the operons in the MPRA dataset (Fig. 5A). We classify sites as promoters only if they have been previously mapped as such; otherwise, we label them as activator sites. Additionally, binding sites located within 5–6 bp of each other are reported as a single site for a more conservative assessment.

**Figure 5.**
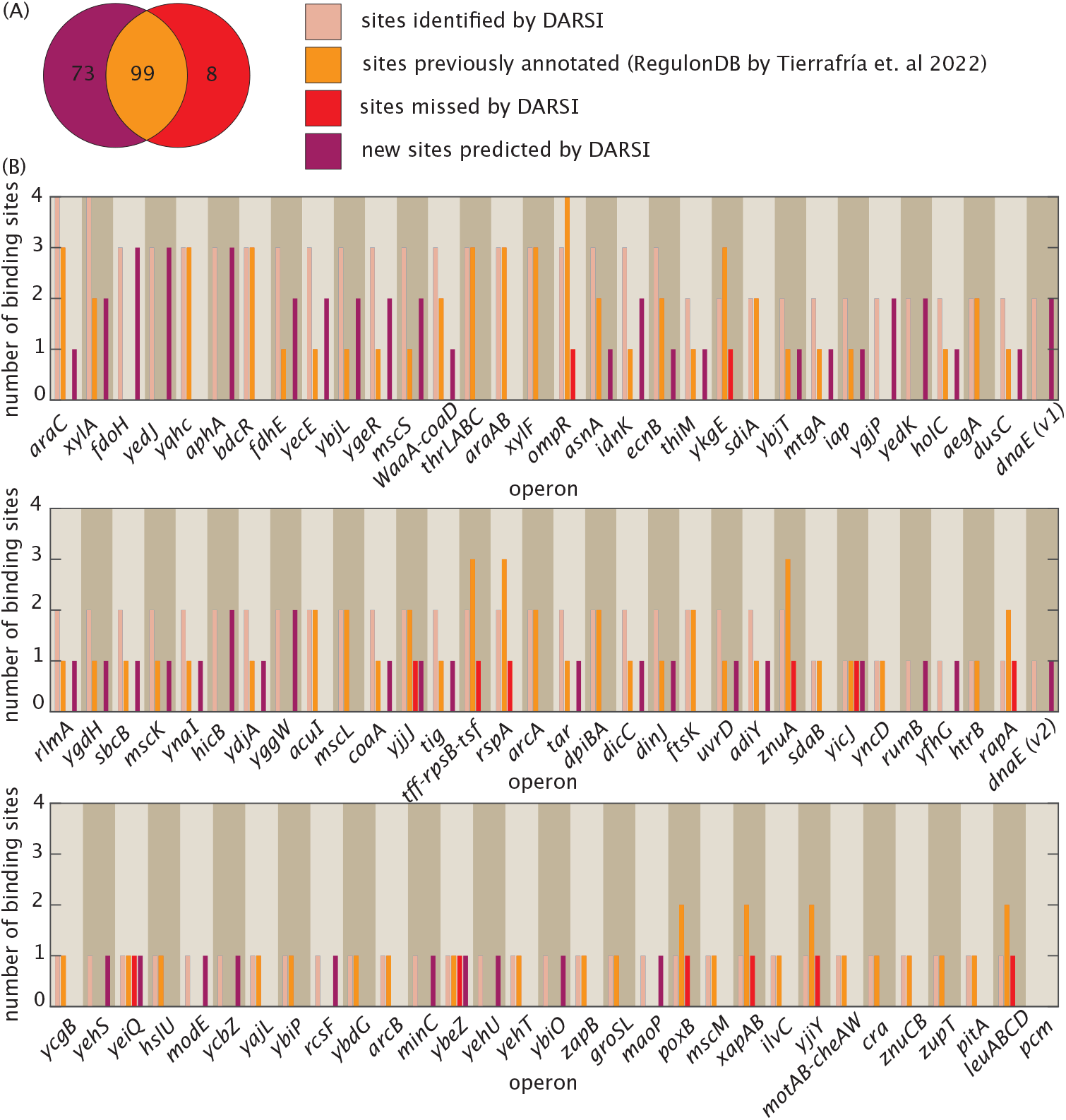
Benchmarking binding sites sites identified by DARSI against curated RegulonDB dataset. **(A)** Venn diagram showing the number of binding sites—both transcription factor binding sites and promoters—identified by DARSI and how that number is compared to the previously known sites. DARSI identified a total of 172 binding sites across all 95 operons, capturing 88.4% of previously known sites documented in published and curated databases (***Tierrafría et al., 2022***), and predicted the existence of 73 new binding sites. **(B)** Bar plots illustrating the number of sites uncovered by DARSI, the number of sites previously annotated in RegulonDB ***Tierrafría et al. (2022***), and the newly identified sites and the sites missed by DARSI across all 95 operons analyzed in this study.

Figure 5B provides a detailed summary of the binding sites identified by DARSI, the missed binding sites, and the newly predicted hypothetical sites for each operon, alongside annotations from the RegulonDB database (***Tierrafría et al., 2022***). Notably, DARSI failed to identify 13 previously annotated sites in RegulonDB for the *ykgE, yicJ, rapA, yeiQ, ybeZ, yjjJ, tff-rpsB-tsf, poxB, rspA, ompR, yjiY, znuA*, and *leuABCD* operons. These missed sites were primarily located within regulatory architectures containing multiple binding sites, such as the *ykgE* and *ompR* operons. Examples of regulatory architectures inferred through DARSI are presented in Figure S3, with comprehensive visualizations for all operons accessible via the GitHub repository. Further, detailed information, including the sequences of each binding site, their genomic coordinates, strand orientation, and prior annotations, is provided in the supplementary table, available for download on the GitHub repository.

## Discussion

MPRAs have become a fundamental experimental tool in the high-throughput dissection of the regulatory genome. The data stemming from these experiments has been matched by an increasingly sophisticated suite of approaches to extract as much information as possible. However, it is clear that there is still much room for improvement. For instance, conventional approaches for finding transcription factor binding sites and promoters such as mutual information rely on local measures and assume independence between base pairs (***Ireland et al., 2020***).

This study highlights the potential of breaking free from the base pair independence assumption and accounting for possible interactions between distant base pairs in a regulatory sequence when finding binding sites within that sequence. In particular, we explored whether the convolutional neural network-based framework embodied in DARSI could enhance the identification of regulatory binding sites and improve our understanding of their roles in dictating gene expression.

To identify binding sites within regulatory sequences using MPRA data, we first demonstrated that DARSI accurately predicts gene expression levels directly—albeit discretely—from raw regulatory sequences. Importantly, this predictive power is achieved without any underlying assumptions about the physical mechanisms and regulatory grammar dictating gene expression. Building on this foundation, we leveraged saliency maps to highlight regions of high information density that drive model predictions to infer locations of transcription factor binding sites and promoters, which are indistinguishable in this approach. While saliency maps provide valuable insights, it is important to acknowledge their limitations, particularly when applied to discrete variables like base pair identity, as this can introduce challenges in interpretation due to the underlying reliance on derivatives (***Kim et al., 2019***).

Trained DARSI models identified over 170 binding sites across all 95 operons, including 73 previously unannotated sites and 99 previously mapped sites, accounting for approximately 90% of previously annotated sites in curated databases. These findings establish DARSI, and convolutional neural networks more in general, as a valuable platform for advancing experimental studies of regulatory architectures. For instance, large *in silico* sequence libraries could be generated to complement and refine *in vivo* experiments, facilitating the design of regulatory sequences tailored for specific expression levels (***de Almeida et al., 2024***; ***Rafi et al., 2024***).

To better understand the effectiveness and limitations of DARSI, we benchmarked its predictions of gene expression sensitivity to mutations against those obtained using traditional mutual information approaches (***Ireland et al., 2020***; ***Kinney et al., 2010***). Figure S4 presents these comparisons for three representative operons. This comparison highlights how peaks are not always identified by both measures of MRPA data, and how those peaks can be slightly displaced and broader in DARSI with respect to those identified by mutual information. A detailed examination of these two measures—highlighting the potential advantages and challenges of DARSI with respect to mutual information—is provided in section “Comparing the DARSI and Mutual Information Approaches” of the Supplementary Information. While, as discussed in that section, the “black box” nature of machine learning models makes it challenging to dissect the source of these difference, the ultimate proof of the usefulness of DARSI and how it compares against well-established approaches will have to stem from future experiments aimed at validating hypothesized binding sites.

It is important to note that while DARSI effectively identifies binding sites, it does not predict the specific transcription factors that bind to these hypothetical sites. Addressing this limitation— as well as elucidating their molecular mechanisms and characterizing their biophysical properties such as binding affinity—remains a significant challenge that will require the integration of additional computational approaches and experimental validation (***Belliveau et al., 2018***; ***Ireland et al., 2020***; ***Pan et al., 2024***). These efforts are essential for advancing our understanding of transcriptional regulation and for improving the utility of predictive models like DARSI in functional genomics.

Although this study focused on bacterial regulatory sequences, the DARSI framework is broadly applicable to differential expression datasets across both bacterial and eukaryotic systems. Beyond gene expression and regulatory sequence activity, DARSI could be adapted for other phenotypic analyses, uncovering causal links and axes of variation in sequence-to-phenotype relationships. For instance, the DARSI methodology could be used to predict protein properties such as solubility, hydrophobic surface composition (***Sato et al., 2023***), or binding affinity (***Littmann et al., 2021***; ***Jones et al., 2021***), given appropriate training datasets.

Importantly, as with other machine learning approaches, the efficacy of DARSI depends heavily on the quality and scale of its training data. Advances in mutagenesis technologies leading to larger MPRA datasets with higher number of variants and broader coverage promise to further amplify the utility of frameworks like DARSI, opening new avenues for precision in computational biology. In particular, our lab envisions exploring DARSI in the context of new mutagenesis technologies that will make it possible to implement MPRAs in multicellular organisms (***Falo-Sanjuan et al., 2024***).

## Materials and Methods

### MPRA Dataset from *E. coli*

The dataset used throughout this article was generated through the work by ***Ireland et al. (2020***). Here, as shown diagrammatically in Figure 1A, a 160-bp long region around the transcription start site of 114 operons in *E. coli* were randomly mutated at a 10% rate (i.e., each base pair along the sequence had a 10% chance of being mutated from its wild-type base to any of its three alternative bases). This library was then cloned into plasmids driving the expression of a reporter gene (Fig. 1B,C). Plasmid libraries were transformed into cells and grown in various growth conditions (though in this work we only focus on bacteria grown in LB). The expression from each operon variant was measured by sequencing (Fig. 1C).

To normalize for the variation in copy number for each reporter construct, DNA counts of the barcode were also included in the table and were used to normalize the expression counts as discussed in the main text. Processed dataset, therefore, provides both the sequence of the regulatory region and the normalized expression count of the gene regulated by that sequence (Fig. 1E).

In this study, we only considered operons with enough sequence variants to ensure that, on average, each base pair was mutated in at least 100 variants (i.e., 100x coverage). Given a 10% mutation rate, this corresponds to a minimum of ∼ 1, 000 variants. Examples of some of these sequence variants within this dataset for the *yqhC* operon are provided in Table S1.

### RNA-seq Raw Data Processing

The sequencing datasets used in this work are deposited in the SRA database as PRJNA599253 and PRJNA603368. Code for sequence processing is provided in the Github repository together with example datasets and Jupyter Notebooks that display how to use the data to generate, for example, Table S1. Here, we give a brief description of the process.

Random barcodes were cloned between the promoter and the reporter gene in order to identify the promoter variant through the RNA reads, as well as provide multiple distinct data points that reduce possible bias introduced by barcodes. In an initial sequencing run the promoter sequence and barcodes were sequenced simultaneously to obtain a map that links a regulatory region variant with the corresponding barcode. Pair end reads were merged, quality filtered, and filtered for read length using “fastp” (***Chen et al., 2018***). Promoter sequence and barcode were extracted from each read and the number of occurrences of each barcode and promoter combination counted. A promoter variant can have multiple barcodes associated with it, however, a barcode has to be unique. If a barcode was observed for multiple promoter variants, the barcode was then removed. Additionally, combinations with less than 3 reads were removed due to the possibility of sequencing errors.

In Reg-Seq, the promoter library is grown in various growth conditions to assess a variety of regulatory conditions. For the purpose of this paper, we take one of these growth conditions: growth in LB. From each culture both RNA and DNA (plasmids) are extracted. Using specific primers, the reporter gene mRNA, including the barcode, is reverse transcribed and amplified to generate cDNA and measure gene expression. Barcodes are also amplified from plasmids using PCR in order to count the number of plasmids present with a specific regulatory sequence. Sequencing adapters are added by another PCR and both barcodes obtained from cDNA and plasmid DNA are sequenced. Reads are trimmed and quality filtered using ‘fastp’ (***Chen et al., 2018***). The occurrence of each barcode is counted in the RNA-Seq and DNA-Seq datasets. Finally, using the results from the initial sequencing run, each corresponding promoter variant is identified through its barcode.

### One-Hot Encoding

Every 160 bp long regulatory sequence from the MPRA dataset is converted to a two-dimensional binary image *A* ∈ ℝ^4×160^, where each entry of the matrix *A*_*i*,*j*_ takes the form

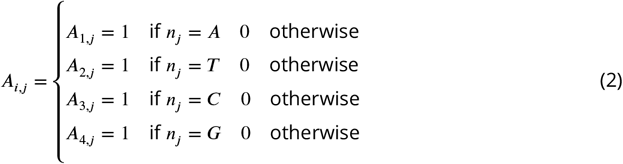

Here, *n*_*j*_ is the *j*^th^ nucleotide in each of the sequences. Figure 2D shows an example of these binary images generated for the regulatory sequences for the *yqhC* operon.

### RNA Count Labeling

The RNA count corresponding to each sequence variant reports on the gene expression level driven by that mutated regulatory region. These values are normalized by dividing the RNA count by the DNA copy number count to ensure that variability in the normalized RNA count is not due to variability in the plasmid copy number.

To bin the normalized expression counts, we developed a binning algorithm to categorize sequence variants into three discrete groups based on their normalized RNA counts: (1) sequences that resulted in zero gene expression, (2) sequences that resulted in low gene expression levels, and (3) sequences that resulted in high gene expression levels. The pipeline first bins all the zero expression counts to the zero expression bin. It then automatically determines the best threshold for separating the remaining gene expression data into the low and high gene expression bins based on their log(normalized mRNA count). To make this possible, a t-test is conducted on the difference of the mean gene expression of the low and high expression bin, leading to a separation of data that minimizes the p-value between the bins.

The vector 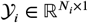 encodes for the expression bins associated with each of the *N*_*i*_ sequence variants for the operon *i* in the dataset. Each sequence variant for a given operon *i* is given a label from a set *K*_*i*_ = {1, 2, 3} where 1, 2, and 3 denote the zero, low, and high expression bins respectively. The algorithm we implemented attempts to partition the vector 𝒴 _*i*_ into three bins such that the p-value associated with a t-test conducted between each pair of bins is minimized (***Mann and Whitney, 1947***; ***Fay and Proschan, 2010***). The iterative algorithm used to bin the normalized mRNA counts is given in Algorithm 1.

#### Algorithm 1 RNA count binning algorithm.

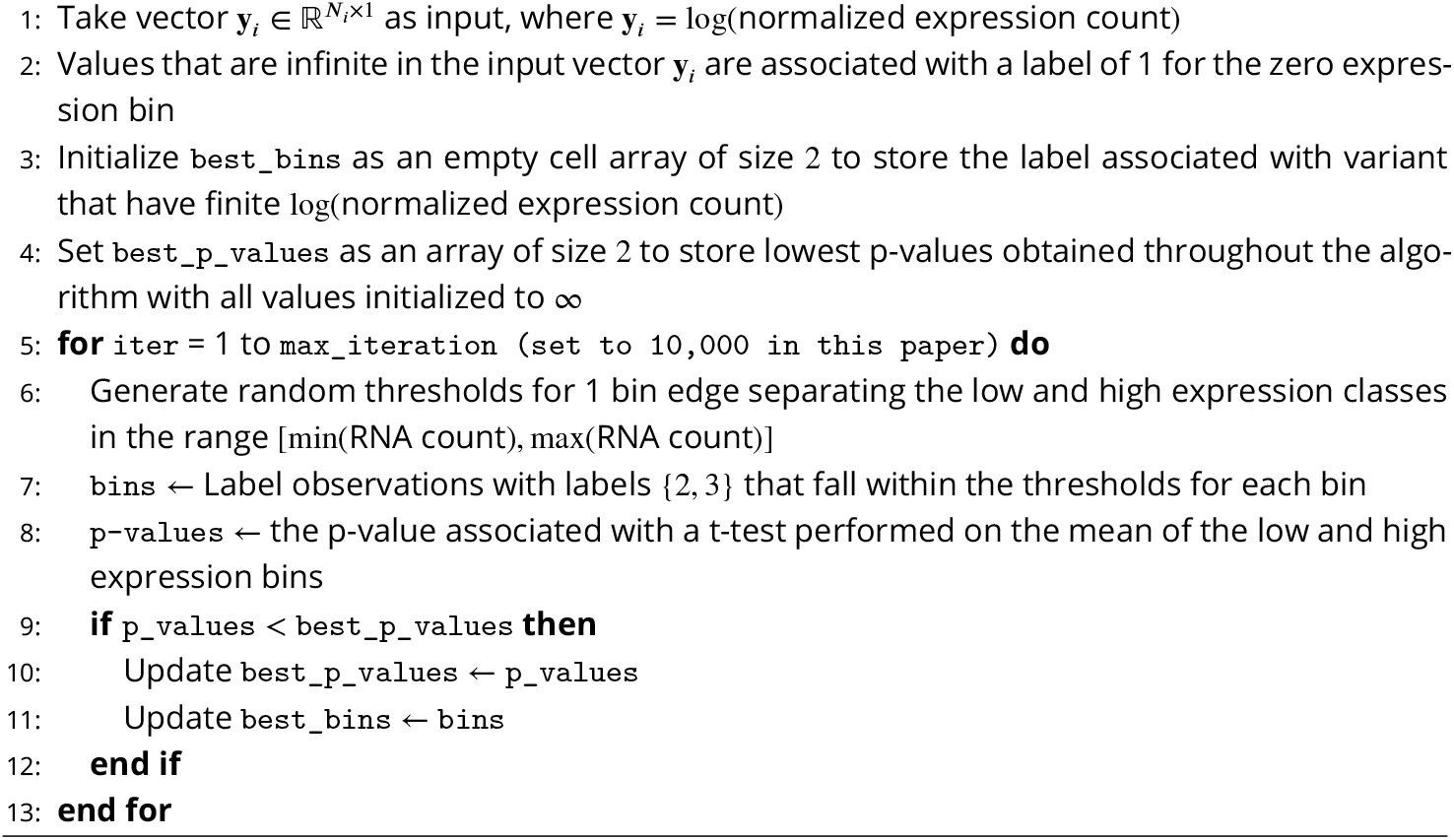

### Saliency Maps

Trained DARSI models demonstrate the capability to predict operon expression levels directly from their nucleotide sequences. Beyond prediction, these models can be utilized to identify potential binding sites within regulatory sequences. This is achieved by analyzing the derivative of the network’s loss function (described in detail below) with respect to the input sequence. By computing these gradients, the nucleotides most critical to the model’s predictive understanding of gene expression can be identified.

The performance of the network is quantified using a cross-entropy loss function defined as

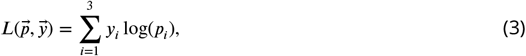

where, 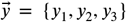 denotes the ground truth label vector for each variant of a given operon. For instance, a variant assigned to the zero expression bin is represented as 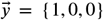. Similarly, 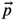 represents the probability vector predicted by the network for the same sequence. This three-dimensional vector specifies the predicted probabilities of the sequence belonging to each expression bin, as determined by the model.

This loss function measures the discrepancy between the network’s predictions and the ground truth, serving as an indicator of model accuracy. To assess the sensitivity of classification outputs to perturbations in nucleotide sequences, we leverage the gradient-weighted class activation mapping (Grad-CAM) approach (***Kudo et al., 1999***; ***Vinogradova et al., 2020***; ***Selvaraju et al., 2017***). Grad-CAM computes the gradient of a selected, differentiable output—such as the cross-entropy loss—with respect to neurons or nodes in a specified layer of the network, typically a convolutional layer. For a toy example demonstrating the computation of saliency maps through backpropagation, please refer to the section “Example of Saliency Map Computation Using Backpropagation” in the Supplementary Information.

This method allows for the visualization of features critical to the model’s predictions by back-propagating the gradients through the network and overlaying them on the input sequence. The resulting gradient map highlights the pixels within the input image (corresponding to nucleotides within the sequence) that significantly contribute to the network’s decision-making process. By identifying these key features, Grad-CAM enhances the interpretability of deep learning models and provides insights into the regulatory architecture underlying gene expression (***Selvaraju et al., 2017***; ***Kudo et al., 1999***; ***Vinogradova et al., 2020***).

For a two-dimensional image classifier such as DARSI, the saliency score for any given channel in a convolutional layer with *k* channels is computed by

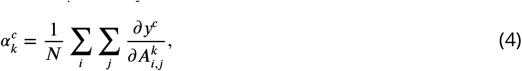

where *y*_*c*_ is the predicted posterior for the bin *c*, 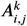 is the pixel located at (*i, j*) position of the *k*^th^ channel of the chosen convolutional layer, and *N* is the total number of pixels (***Selvaraju et al., 2017***).

Equation 4 generates a weighting score for every channel *k* within a convolutional layer. In order to plot this score as a heatmap for any given sequence, similar to the ones shown in Figure 4C and D, a weighted-average mask *U* is computed such that

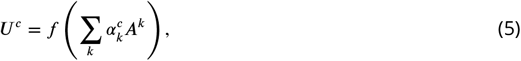

where *f* represents a non-linear activation function such as the rectified linear unit function ReLU *f* = max(0, *x*) (***Selvaraju et al., 2017***). Algorithm 2 summarizes the steps that are taken to generate these saliency maps for each of the genes within the expression shift dataset.

### Binding Site Identification

As discussed in Algorithm 2 above, the saliency vector 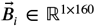 computed for a given operon *i* captures the saliency of each regulatory sequence. The vector 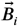 is normalized about its mean and

#### Algorithm 2 Saliency map generation.

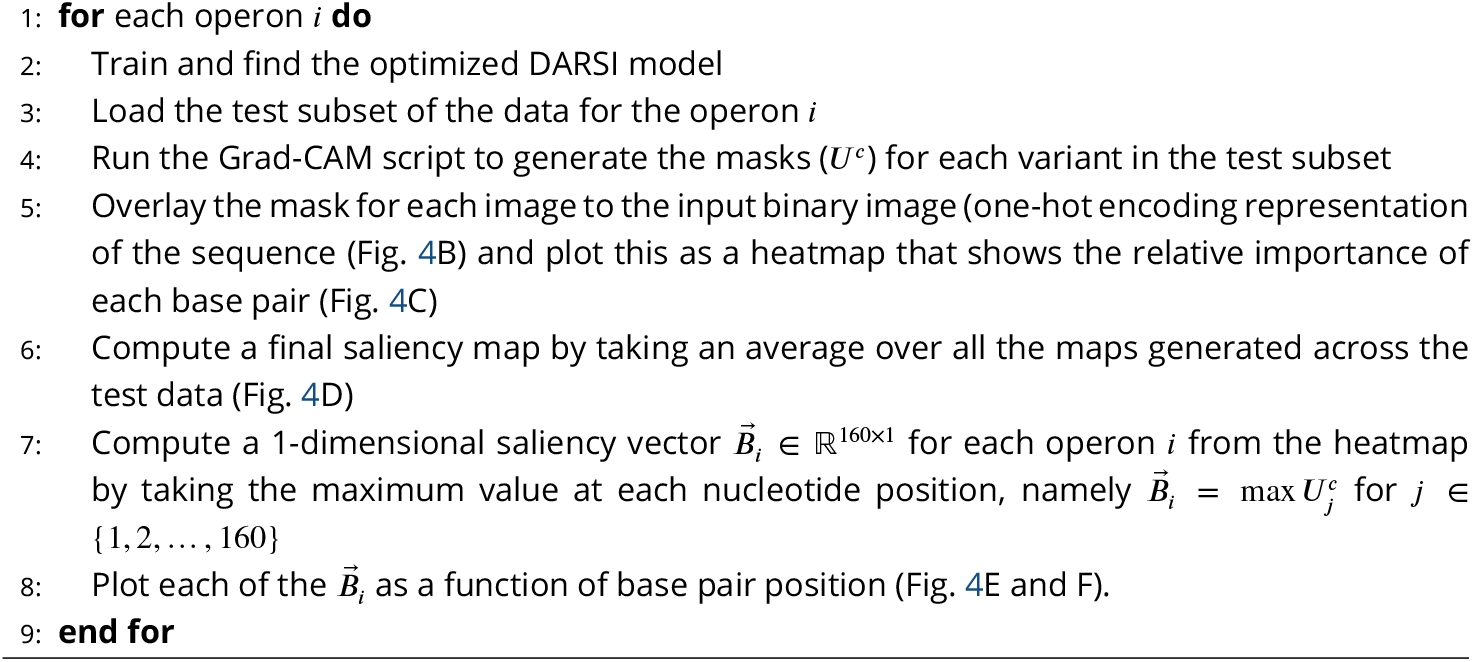

standard deviation, namely

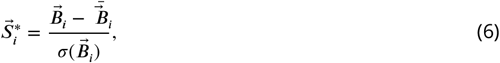

where 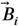 and 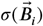 are the mean and standard deviation of the vector 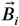, respectively.

The vector 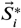 represents a difference from the mean in sensitivity of expression level to mutation at any given position *j*. Therefore we assume that this vector is proportional to the derivative of the dissociation constant with respect to that nucleotide, or more formally

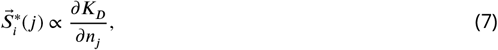

where *K*_*D*_ and *n*_*j*_ are the dissociation constant and nucleotide at position *j*, respectively.

Finally, we used this proportionality to estimate the probability of occupancy *P* (*j*)_*i*_ as

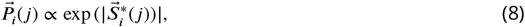

where |·| denotes the absolute value. The use of absolute values is necessary due to prior normalization of the vector 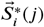, which ensures that both strongly negative and strongly positive normalized values contribute to the probability estimate. This adjustment is crucial to account for regions associated with activators (negative values) and repressors or other functional elements (positive values), ensuring a strong signal is captured in both cases.

The probability was computed for every position *j* for every operon *i* and was plotted as bar charts to show the expression shift (Fig. 4E). The peaks in probability that were more than one standard deviation from the mean were selected (Fig. 4F). These filtered peaks were then passed through a secondary filter to select only regions where the length of a continuous region of repression or activation (i.e. the predicted binding site) is more than 10 bp long—the minimum length of the binding sites in *E. coli* (***Stewart et al., 2012***; ***Rydenfelt et al., 2015***; ***Ruths and Nakhleh, 2013***). The process for generating filtered expression shift plots is shown in Algorithm 3.

### The DARSI Pipeline Repository

To enhance the accessibility and usability of DARSI for gene expression prediction and related applications, we have designed the implementation with a modular architecture. Each *MATLAB* script operates independently, accompanied by comprehensive documentation for ease of understanding. A master script is also provided to sequentially execute the necessary functions, offering detailed guidance on processing raw RNA-Seq data and training a DARSI model.

#### Algorithm 3 Binding sites identification.

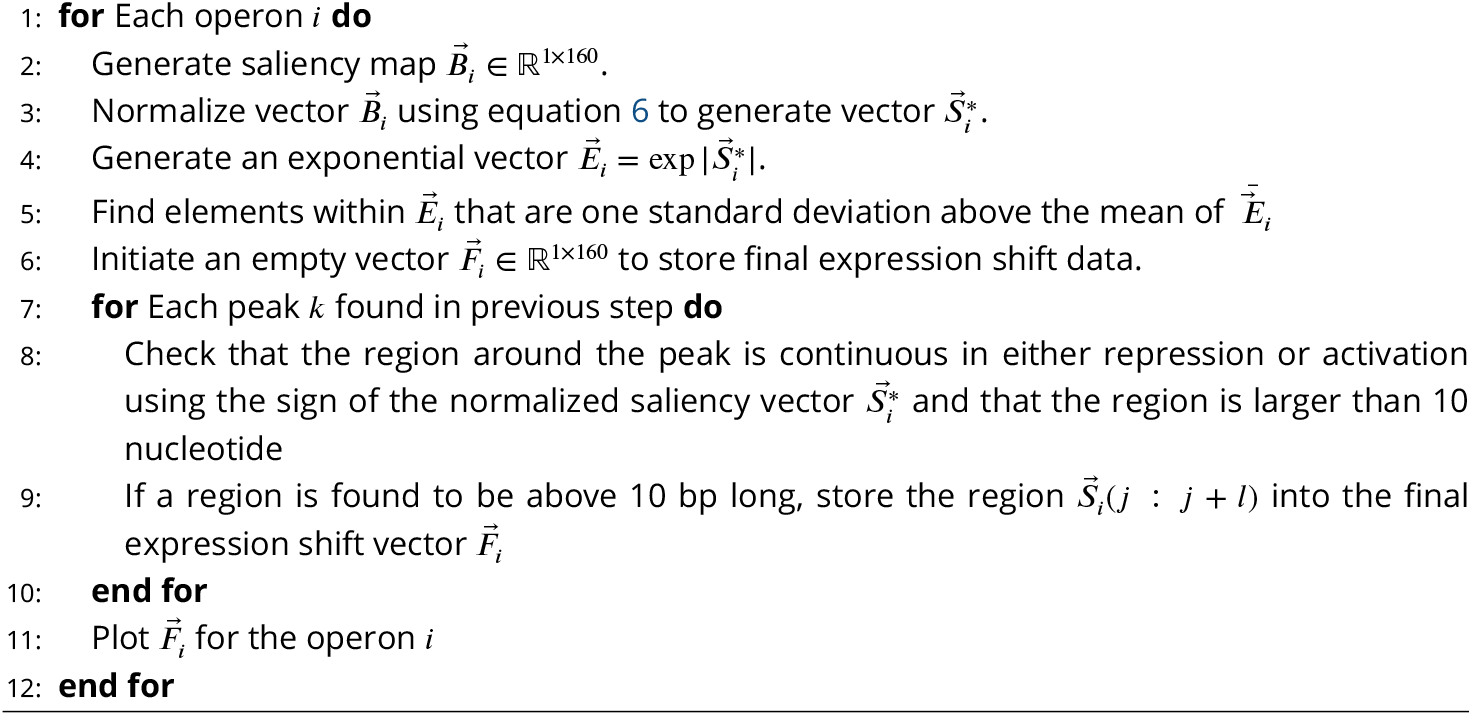

The complete set of scripts is available in a dedicated GitHub repository. The repository includes modules for processing raw RNA-Seq data, generating expression shift datasets, training DARSI models, performing cross-validation, evaluating model performance, and identifying binding sites. Additionally, all data used to train the DARSI model, along with outputs such as saliency maps, confusion matrices, and expression shift plots, are made available in the repository.

## Acknowledgments

We would like to thank Rob Philips, and Julia Falo-Sanjuan for comments on the manuscript. H.G.G. was supported by NIH R01 Awards R01GM139913 and R01GM152815, by the Koret-UC Berkeley-Tel Aviv University Initiative in Computational Biology and Bioinformatics, and by a Winkler Scholar Faculty Award. H.G.G. is also a Chan Zuckerberg Biohub Investigator (Biohub–San Francisco). A.K. was supported by Phyllis B. Blair Graduate Fellowship from the University of California, Berkeley.

## Supplementary Information

### Supplemental methods

#### MPRA Dataset Example

The raw RNA-Seq data from *E. coli* cultures is processed as described in the “RNA-seq Raw Data Processing” section of the Materials and Methods. The processed data is tabulated to form what is referred to as the MPRA dataset throughout this work. Table S1 shows a few rows of this type of data for the illustrative *yqhC* operon.

**Table S1.**
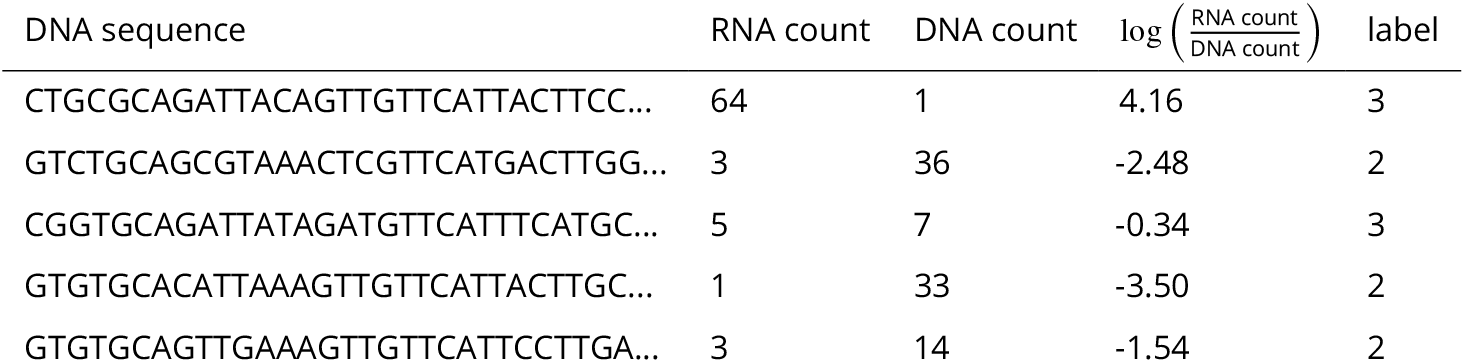
Illustrative example of the differential expression dataset used throughout this work. This table features processed MPRA data for the *yqhC* operon. Each row represents a uniquely mutated 160 bp-long promoter sequence. The dataset includes the following columns: 1) DNA Sequence: The 160 bp-long DNA sequence of the mutated promoter. 2) RNA count: The measured expression level of the reporter gene, as quantified by RNA-Seq. This reflects the transcriptional activity associated with the promoter sequence. 3) DNA count: The count of DNA barcodes corresponding to the copy number of each sequence in the library. This serves as a measure of the copy number of the plasmid containing each regulatory sequence. 4) 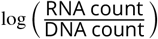: A normalized measure of expression, calculated by dividing the RNA count by the DNA count and taking the logarithm of the result. This normalization accounts for variations in sequence abundance and enables direct comparison of transcriptional activities across sequences. 5) Label: A discretized classification assigned to each sequence, derived from binning the normalized log-expression values into categories based on a binning algorithm (1: zero expression bin, 2: low expression bin, 3: high expression bin) as described in the “RNA Count Labeling” section of the Materials and Methods. This table format is applied consistently across all operons analyzed in this study, allowing for a systematic comparison of promoter sequence variants and their transcriptional activities.

#### DARSI Architecture and Training

The convolutional layers used in DARSI have filters spanning a 5-bp range to capture local sequence patterns and nucleotide interactions. While these filters operate on short sequence windows, stacking multiple convolutional layers extends the effective receptive field, allowing the network to model higher-order interactions and long-range dependencies across the regulatory sequences. In principle, this architecture enables DARSI to detect complex regulatory features, such as distant transcription factor binding site interactions. However, the inclusion of additional convolutional layers increases model complexity, which can lead to overfitting.

To determine the optimal architecture, we utilized data from the 10 operons with the largest number of variants (*leuABCD, rumB, zupT, yncD, uvrD, mscK, ftsK, yqhC, groSL*, and *xylA*). We systematically varied the number of convolutional layers and the number of filters within each layer, evaluating training and validation accuracy to balance model complexity and generalization. Figure S1A depicts the changes in training and validation accuracies as the number of convolutional layers increased from 2 to 6, with each layer comprising 16 channels. The results indicate that, while training accuracy increases monotonically, validation accuracy decreases monotonically, signaling overfitting. This suggests that the optimal number of layers lies between 2 and 3. To mitigate potential overfitting when training the network on smaller datasets for other operons, we selected 2 as the optimal number of layers. With the number of layers fixed, we further examined the impact of the number of channels per layer on training and validation accuracies. Figure S1B illustrates the average training and validation accuracies across the same 10 operons as the number of channels per layer is varied. The optimal number of channels was determined to be 32, as this corresponds to the maximum validation accuracy. Thus, the final architecture we converged on, with 2 convolutional layers and 32 filter counts, achieved comparable training and validation accuracies, minimizing overfitting while preserving predictive power.

The optimal architecture for DARSI network comprises 12 hidden layers and was designed, trained, and evaluated using the *Matlab* Deep Learning Toolbox (***MathWorks, 2022***). The network was trained using stochastic gradient descent with an initial learning rate of 0.001 (***Bottou, 1998***; ***Sra et al., 2011***; ***Ruder, 2017***). Training was performed for a maximum of 20 epochs with a minibatch size of 32, and the learning rate was reduced by 20% every 5 epochs. The training dataset was shuffled at the start of each epoch to improve generalization. Table S2 outlines the layers of the optimized architecture, along with their descriptions, dimensions, and learnable parameters.

#### Example of Saliency Map Computation Using Backpropagation

In neural networks, computing the derivative of the output with respect to the input involves propagating gradients backward through the network—a process known as backpropagation. This task becomes increasingly complex as the number of hidden layers grows and the architecture incorporates advanced components such as convolutional, pooling, and activation layers. To provide clarity and intuition about this process, we include a toy example in this section, illustrating how backpropagation operates specifically through convolutional layers. This example is intended to demystify the mechanics of gradient computation and offer a simplified yet instructive view of how saliency maps are generated.

**Figure S1.**
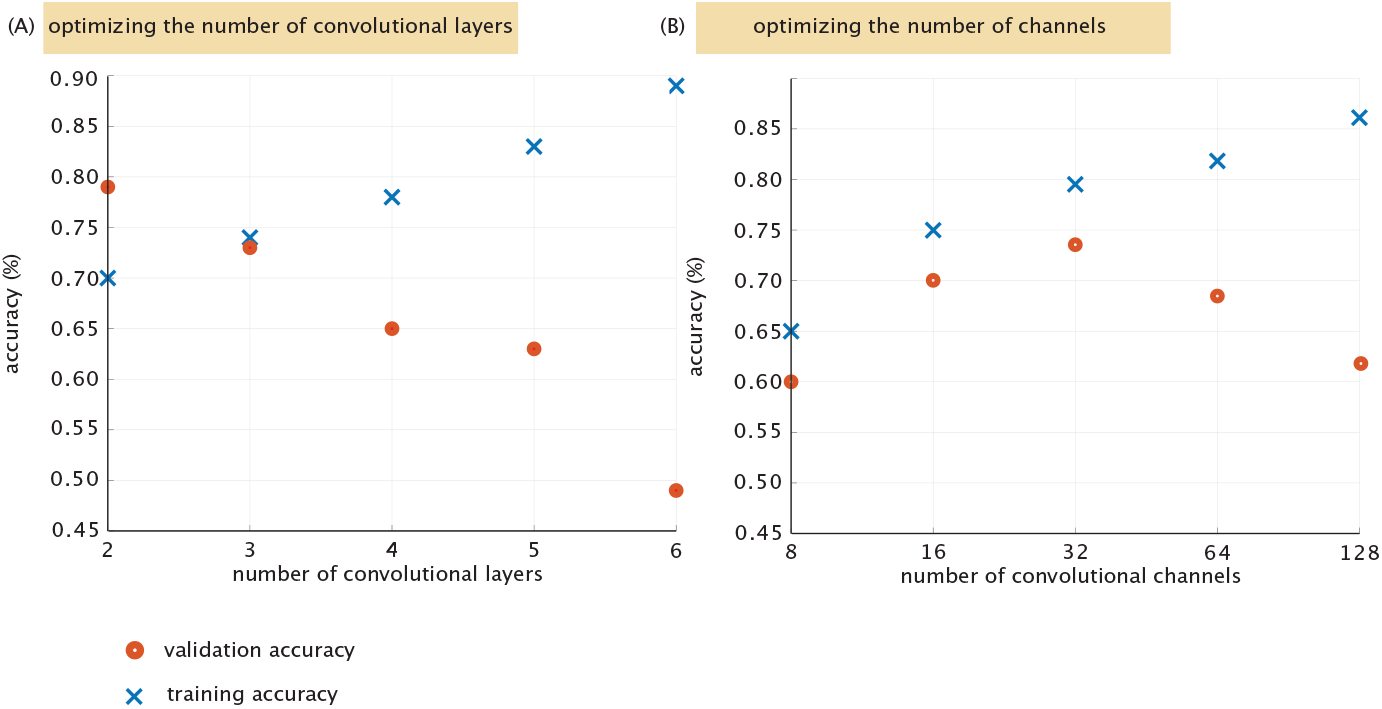
DARSI network architecture optimization. **(A)** The average training and validation accuracy for the 10 operons, with the largest number of variants, used for network optimization (*leuABCD, rumB, zupT, yncD, uvrD, mscK, ftsK, yqhC, groSL* and *xylA*) are plotted against the number of convolutional layers (given 16 channels per layer). **(B)** Given an optimum of 2 convolutional layers, we assay network training and validation accuracy as a function of the number of channels within each convolutional layer. The optimal architecture was selected to achieve comparable or higher validation accuracies (red dots) relative to training accuracies (blue crosses). This criterion ensures a balance between model complexity and generalization, favoring architectures that capture the underlying patterns in the data effectively while minimizing the risk of overfitting. By prioritizing validation performance over excessive improvements in training accuracy, the selected architecture demonstrates robust generalization across diverse datasets.

**Table S2.** Detailed description of all 12 layers of the optimized DARSI architecture along with their description, dimensions, and number of learnable parameters.

| Layer name | Type | Dimension | Learnable properties | Number of learnable parameters |
| --- | --- | --- | --- | --- |
| Sequence Input | Image input | 4x160x1x1 | - | 0 |
| Conv_1 | Convolution | 4x5x1x32<br>stride [1 1]<br>padding same | Weights: 4x5x1x32<br>Biases: 1x1x32 | 672 |
| Batchnorm_1 | Batch Normalization | 32 channels | Offset: 1x1x32<br>Scale: 1x1x32 | 64 |
| Relu_1 | ReLU | N/A | - | 0 |
| Maxpool_1 | Max Pooling | 1x2<br>stride [2 2]<br>padding [0 0 0 0] | - | 0 |
| Conv_2 | Convolution | 1x5x32x64<br>stride [1 1]<br>padding same | Weights: 1x5x32x64<br>Biases: 1x1x64 | 10,304 |
| Batchnorm_2 | Batch Normalization | 64 channels | Offset: 1x1x64<br>Scale: 1x1x64 | 128 |
| Relu_2 | ReLU | N/A | - | 0 |
| Maxpool_2 | Max Pooling | 1x2<br>stride [2 2]<br>padding [0 0 0 0] | - | 0 |
| Fc | Fully Connected | 1x1x3 | Weights: 3x1<br>Biases: 3x1 | 6 |
| Softmax | Softmax | 1x1x3 | - | 0 |
| Classoutput | Classification Output | 1x1x3 | - | 0 |

Consider the sequence shown on the left of Figure S2A. To compute the derivative of the network’s loss function, 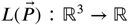, with respect to each position in the input sequence *x*_*i*_, we use the cross-entropy loss function, defined as

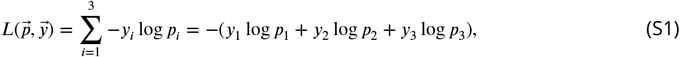

where *p*_*i*_ represents the probability that the sequence belongs to expression bin *i*, and 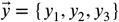 is a binary one-hot encoded vector indicating the ground truth class. In 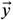, only one element is 1, corresponding to the correct class, while the rest are 0. For this example, assume the sequence belongs to the low-expression class. This results in a ground truth vector of 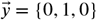.

As shown in Figure S2A, the input sequence is fed into the trained DARSI model to generate the value for 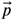. We can, therefore, write this forward pass of the sequence through the network as

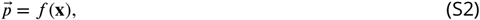

where × represents the input sequence and *f* (*x*) : ℝ^4×160^ → ℝ^3^ denotes all the layers of trained DARSI network, all lumped together in *f* (x).

To compute the saliency map for the sequence image x, we calculate the gradient of the loss function with respect to each position *x*_*ij*_ in the input sequence by applying the chain rule iteratively from the output layer to the input layer of the network such that

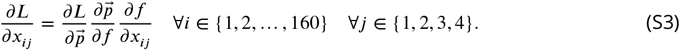

Because *f* (x) consists of multiple layers and transformations, computing this derivative requires an iterative approach. Starting from the loss function, we iteratively propagate gradients backward through the network using the chain rule of differentiation, layer by layer, until the input is reached. This procedure, known as backpropagation, emphasizes the reverse traversal of layers to compute the necessary gradients.

To illustrate the computation of gradients through a convolutional layer, we present a detailed example. Consider a matrix *A* ∈ ℝ^2×2^ as the input to a two-dimensional convolutional layer to which a single filter *W* ∈ ℝ^1×2^ is applied, resulting in an output *y* = *f* (*A*), as depicted in Figure S2B. This example demonstrates the step-by-step process of propagating gradients through the convolutional layer.

First, the convolution operation is performed using a filter with known parameters *W* = [*w*_1_, *w*_2_], that were obtained in the training process. The resulting output is transformed by applying a non-linearity, in this case, the logarithmic function. The output of the convolution is expressed as

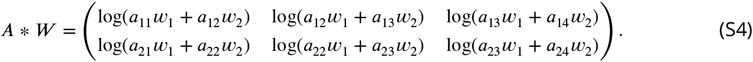

The final output of this convolutional layer is a non-linear transformation *y* = *q*(*A* * *W* ), where *q*(.) denotes the logarithmic activation function. To compute the gradient of the output *y* with respect to a specific input element *a*_*ij*_ , we apply the chain rule of differentiation. For instance, the derivative with respect to *a*_11_ is given by

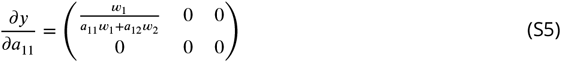

This calculation illustrates how convolutional layers integrate contributions from neighboring elements of the input matrix. For example, the value *a*_12_ contributes to the saliency computed for *a*_11_, highlighting the localized yet interconnected nature of saliency computation in convolutional architectures. This same approach to backpropagation can be used to compute the saliency maps by propagating from the loss function as shown in Figure S2A.

**Figure S2.**
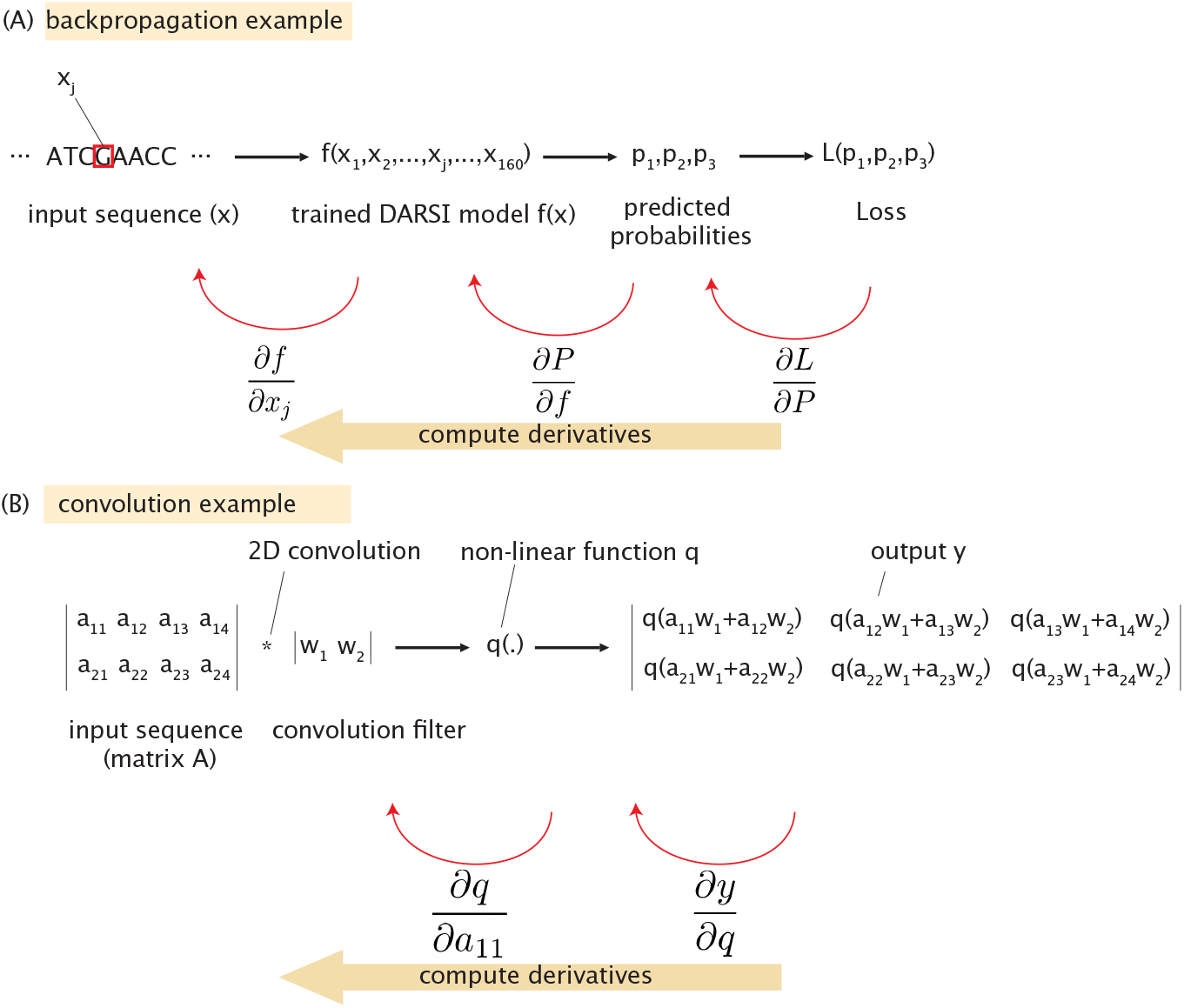
Illustration of the backpropagation process and convolution operation in the DARSI model. **(A)** Example of backpropagation with an input DNA sequence processed by the trained DARSI model *f* (*x*), which predicts probabilities *p*_1_, *p*_2_, *p*_3_. The loss function *L*(*p*_1_, *p*_2_, *p*_3_) is computed using the probability vector 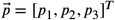. **(B)** Demonstration of a 2D convolution operation applied to a segment of the input sequence using a filter *w*_1_, *w*_2_, followed by a non-linear activation function *q*(*x*) = log *x*.

### Comparing the DARSI and Mutual Information Approaches

Most peaks in DARSI saliency plots correspond to regions of high mutual information reported by ***Ireland et al. (2020***), affirming the capacity of DARSI to detect key regulatory elements. Figure S4 highlights peaks for activating and repressing regions that show strong overlap between the two methods. Specifically, in Figure S4A we show how, for the *dpiBA* operon, DARSI identifies a clear peak which aligns with a region of high mutual information.

While DARSI and mutual information footprints share similarities, there are notable differences between these two measures. For example, in Figure S4B we present a comparison between DARSI and mutual information for the *coaA* operon. The figure shows that, while there is a clear correspondence between peaks reported by DARSI and mutual information, a relative shift of the peaks can be observed. We speculate that these slight positional shifts can occur because the convolutional layers output values are processed by a maximum pooling layer. This layer selects the highest value within a 2 bp window, effectively averaging the signal over small regions and potentially shifting windows by 1 or 2 bp.

The differences between DARSI and mutual information also become obvious in the context of the *yqhC* operon shown in Figure S4C. The figure shows how while some peaks are only identified through DARSI, some other peaks are only present through the mutual information description. Comparative plots, similar to Figure S4, for all 95 operons in this study can be found in the GitHub repository.

Overall, Figure S4 suggests that peaks generated by DARSI tend to be broader, which may reflect either a biologically meaningful characteristic—such as broader peaks capturing actual regulatory sites—or a consequence of information diffusion through the network’s convolutional layers. Further, DARSI identifies more continuous regions of activation and repression, characterized by smooth and extended stretches of blue or red bars in the saliency plots, whereas mutual information plots often exhibit scattered, discrete regions of activity. These differences may reflect the assumption of independence between base pairs underlying mutual information analysis, or potential overfitting in DARSI’s predictions.

Importantly, given the complexity of the DARSI architecture—and as the case with most neural networks—dissecting the inner workings of the network to explain the observed differences between its outputs and those of conventional methods remains highly challenging and speculative. Indeed, because of the ultimate “black box” nature of DARSI, an important limitation of the saliency maps generated with this network is their lack of direct physical interpretation: they are unitless in contrast to the interpretable, information-theoretic units provided by mutual information (bits). Despite these drawbacks with interpretability, DARSI’s ability to incorporate nucleotide interactions offers a complementary perspective that extends beyond the scope of traditional methods.

## Supplemental Figures

### Gene Expression Sensitivity to Mutation Plots

Examples of gene expression sensitivity plots for three illustrative operons along with cartoon of their inferred regulatory architecture is given in Figure S3. Similar plots for all 95 operons can be found in the [GitHub repository](https://github.com/armankarshenas/DARSI/tree/main/plots/expression_plots).

### Expression Plots Comparison

Figure S4 shows examples of unfiltered expression sensitivity-to-mutation plots generated by DARSI stacked against mutual information plots implemented as discussed in ***Ireland et al. (2020***). Similar comparison plots for all 95 operons can be found in the [GitHub repository](https://github.com/armankarshenas/DARSI/tree/main/plots/comparison_plots).

**Figure S3.**
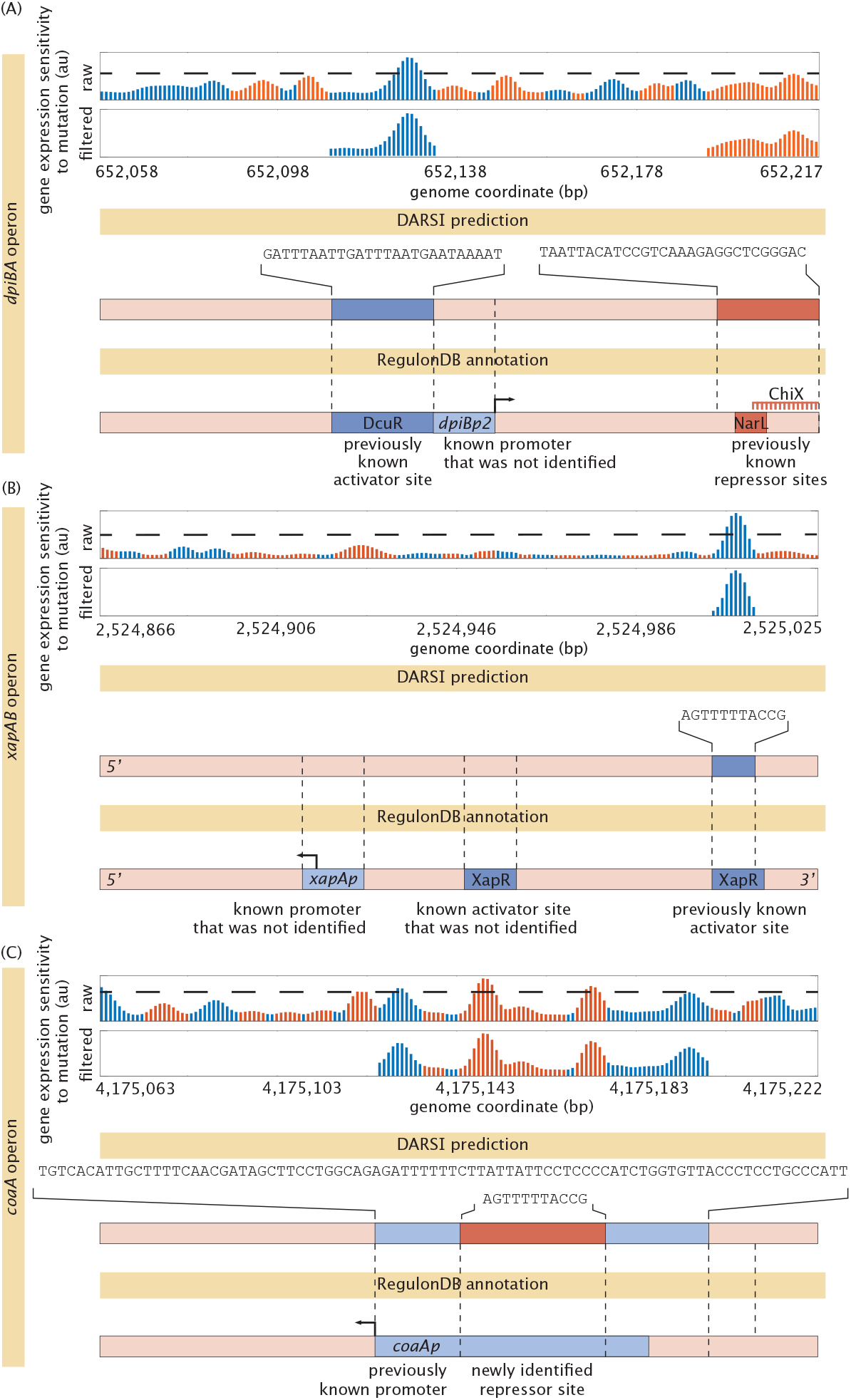
Illustrative examples of expression sensitivity plots. Plots of raw expression sensitivity to mutation are presented for three illustrative operons selected to demonstrate distinct scenarios of the performance of DARSI. Each raw sensitivity plot is accompanied by its filtered version (obtained using the theshhold indicated by the dashed line), which was used to infer the location and type of binding sites (activators vs. repressors). Additionally, regulatory cartoons depict the predicted binding sites, their sequences, and previous annotations based on RegulonDB (***Tierrafría et al., 2022***). It should be noted that the sequences are always presented in the 5’ to 3’ direction regardless of the strand. **(A)** Shows how DARSI successfully identified, as well as missed, previously annotated binding sites in the *dpiBA* operon. **(B)** While DARSI identified an already known site in the *xapAB* operon, it failed to another already known binding site as well as the promoter. **(C)** In the *coaA* operon, DARSI successfully identified the promoter as well as predicted a new repressor site.

**Figure S4.**
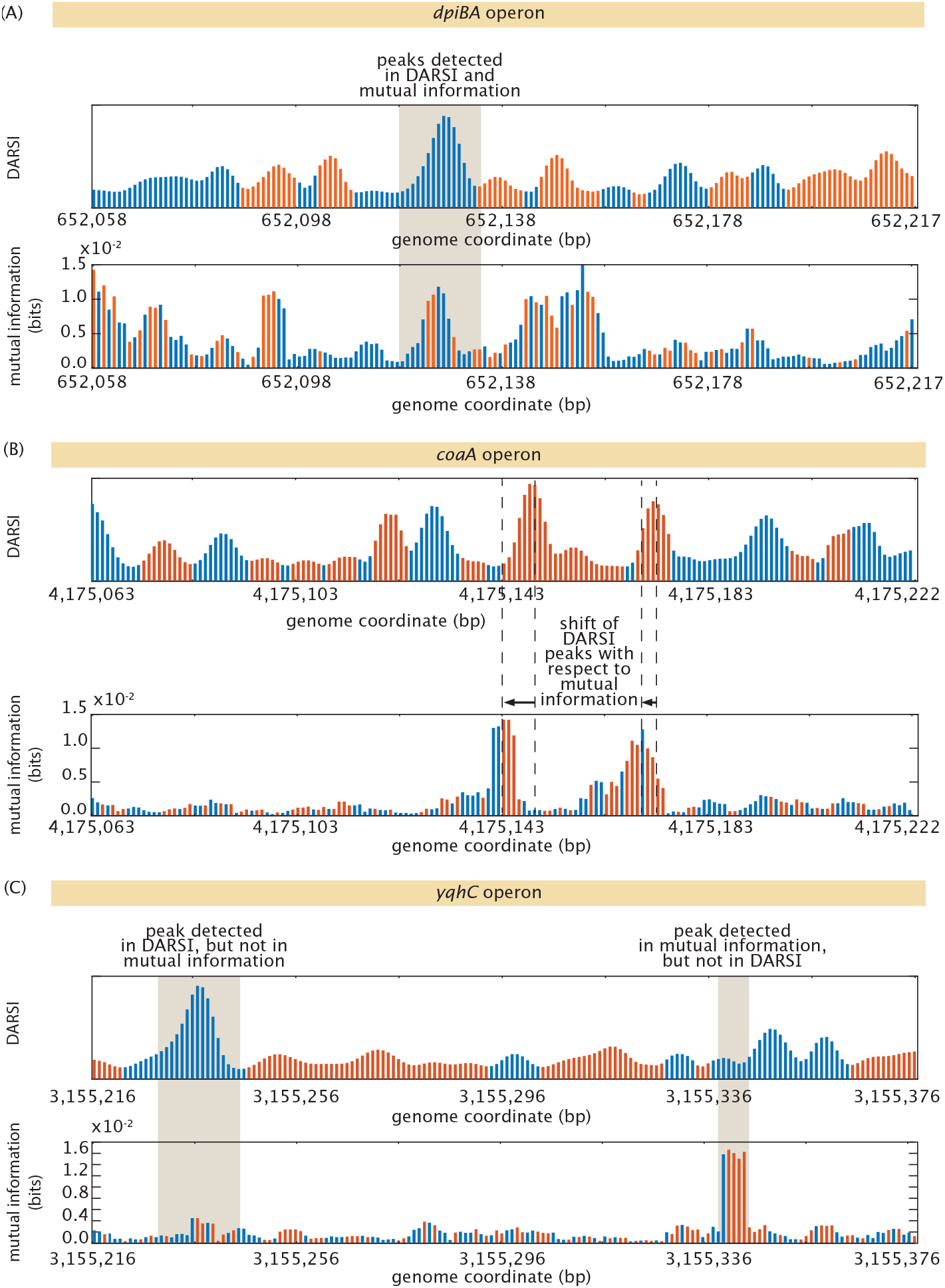
Illustrative DARSI gene expression sensitivity plots compared with mutual information. Gene expression sensitivity-to-mutation plots generated by DARSI and mutual information for three illustrative operons. (A) The *dpiBA* exemplifies an agreement of some peaks detected by DARSI and by mutual information. (B) These peaks, however, can be displaced between these two measures as shown here for the *coaA* operon. (C) Analysis of the data for the *yqhC* operon reveals that peaks can be found by one measure but not the other one.

